# Positive indirect effects more than balance negative direct effects of ungulate grazers on population growth of a grassland herb

**DOI:** 10.1101/2022.07.24.501324

**Authors:** Thomas J. Richards, Michel Thomann, Johan Ehrlén, Jon Ågren

## Abstract

1. Herbivores can affect plant population dynamics both directly because of the damage they inflict, and indirectly by moderating conditions for plant recruitment, competition and other biotic interactions. Still, the relative importance of indirect effects of herbivores on plant population dynamics is poorly known.
2. We quantified direct and indirect effects of ungulate grazers on population growth rate of the short-lived perennial herb *Primula farinosa*, using integral projection models based on demographic data collected over 7 years in exclosures and open control plots at nine grassland sites in southern Sweden. In addition, we explored the mechanisms behind indirect effects with simulations.
3. Grazers had negative direct effects on *P. farinosa* population growth rate, but these were more than balanced by positive indirect effects. The positive indirect effects were mainly linked to improved conditions for plant recruitment. Simulations indicated that indirect effects of ungulate grazers on population growth rate via interactions with pollinators, seed predators, and small herbivores were weak in this system.
4. *Synthesis*. The results demonstrate that a full understanding of the effects of grazing on plant population dynamics requires that both direct and indirect effects are identified and quantified. Plant species vary considerably in their response to shifts in grazing regime. Our study sets an example for how the causes of such variation can be assessed, and thus providing a better understanding of the variable effects of herbivores on plant fitness, abundance and distribution.

## Introduction

Identifying the main factors determining population growth and fitness is key for predictions of responses to environmental change (Morris et al. 2020). Many abiotic and biotic factors may affect population growth rate both directly and indirectly. For example, grazers can affect the vital rates and population dynamics of a given plant species both directly because of the damage they inflict to this species, and indirectly by influencing abiotic conditions and biotic interactions. Indirect effects of grazers on plant vital rates have been identified in both grassland and forest ecosystems (Hobbs 1996; Rooney & Waller 2003). However, similar to other types interactions where indirect effects on individual components of fitness have been demonstrated, the relative importance of direct and indirect effects of grazing intensity on plant population dynamics is poorly understood.

The overall effect of grazers on the population dynamics of plants varies among species as reflected in the many times striking changes in plant community composition when grazers are introduced or removed from an area (Huntly 1991; Olff & Richie 1998; Bakker et al. 2006; Young et al. 2013; Beck et al. 2015; Borer et al. 2015). Plants vary in tolerance to damage, but in general the direct effects herbivory on plant fitness will be negative with the strength of the effect depending on the part of the plant and the timing of damage relative to plant development (Marquis 1992; Gómez 2005; Lehndal & Ågren 2015). In contrast, indirect effects of herbivores can affect the fitness of focal species both positively and negatively. Trampling can create bare patches that facilitate seedling establishment (Collins 1987; Belsky 1992; Bullock et al. 1994; Jutila & Grace 2002), and selective grazing may reduce competition for light and nutrients for plants that escape damage (McNaughton et al. 1997; Veen et al. 2008; Borer et al. 2015; Koerner et al. 2018; Orr et al. 2022). Reduced vegetation height as a result of grazing may in turn affect several other biotic interactions. For example, it may increase pollination success, but also result in increased seed predation of short plants interacting with visually searching pollinators and seed predators (Toräng et al. 2008). Furthermore, reductions in vegetation height and litter accumulation will reduce cover for rodents and other small herbivores and may thus decrease damage by these herbivores (Maclean et al. 2011). The net effect of grazers on population growth of a focal plant species will thus often not be possible to assess without knowing the relative strength of direct negative effects, and positive and negative indirect effects.

Determining the total effects of ungulate grazers on plant population growth rate requires that vital rates are documented across the life cycle of the plant, and population growth estimated in areas with and without grazing (Gómez 2005; Maron & Crone 2005). Direct effects can be assessed by comparing population growth between damaged and undamaged plants (cf. Ehrlén 2003), and indirect effects can be quantified as the difference between total and direct effects. An appropriate assessment of direct and indirect effects of grazers may often require manipulations lasting for several years. First, if exclosures are established in an area with a history of grazing, the estimate of indirect effects of grazing on population growth rate can be expected to grow over time as vegetation recovers from previous grazing and grows taller inside exclosures, and stabilize only after a few years. Second, among-year fluctuations in environmental conditions can be expected to influence the strength of both direct and indirect effects of grazing.

Here, we apply integral projection models to demographic data collected in an exclosure experiment conducted at nine sites over 7 years to quantify total, direct, and indirect effects of ungulate grazers on the population dynamics of the short-lived grassland herb *Primula farinosa*. Previous studies suggest that this species is favoured by moderate disturbance of the surrounding vegetation. First, historical observations indicate that grazing and mowing promote *P. farinosa* population persistence (Lindborg & Ehrlén 2002). Second, ungulate grazers strongly reduce vegetation height in the grasslands of the study area (Thomann et al. 2018), and reduction of the height of surrounding vegetation increased fruit and seed production of *P. farinosa* in a field experiment (Ågren et al. 2006). Finally, grazing and associated reduction in vegetation height and litter accumulation, and the disturbance of the ground cover through trampling should improve conditions for seedling establishment, reduce resource competition from other plant species, and reduce herbivory from rodents. Based on these premises, we tested the following hypotheses: (1) Population growth rate of *P. farinosa* is higher in the presence of ungulate grazers than when ungulates are absent, (2) Direct effects of grazing, in terms of damage to exposed individuals, affects population growth negatively, whereas (3) Indirect effects on population growth rate are both positive and negative. To distinguish between indirect effects of grazers via interactions with pollinators, seed predators, and small herbivores, we estimated the effects of these biotic interactions on vital rates inside and outside exclosures, and tested the specific predictions that grazers by reducing vegetation height positively affect *P. farinosa* population growth rate through increased pollination success and reduced damage from small herbivores, but that grazing also have negative effects by increasing seed predation.

## Materials and methods

### Study system

*Primula farinosa* is a perennial herb with a basal rosette of up to 5 cm in diameter. It is hermaphroditic and self-incompatible, and its flowers are arranged in an umbellate inflorescence on a 0-20cm scape (Hambler & Dixon 2003; Ågren et al. 2013). It has a disjunct distribution in Europe and inhabits moist calcareous grasslands from central Sweden and Scotland in the north to high-altitude areas in Spain, France and Bulgaria in the south. This study was conducted in alvar environments on the island Öland, southeastern Sweden. In this area, *P. farinosa* grows in open semi-natural calcareous grasslands grazed by deer and moose, and more regularly by domestic grazers including cattle, horses and sheep. After a period of low grazing intensity, grazing has increased since the 1990s. In the study area, *P. farinosa* flowers from early May to early June and is mainly pollinated by the solitary bee *Osmia bicolor* and the small butterfly *Pyrgus malvae*. Fruits mature within 6–8 weeks. Fruits can be eaten by larvae of the moth *Falseuncaria ruficiliana*, which develop in the maturing fruit. Fruits attacked by the seed predator produce no seeds. On Öland, *P. farinosa* is dimorphic for scape length, with a long-scaped morph displaying its flowers well above the ground, and a short-scaped morph with its inflorescence located close to the ground (Ehrlén et al. 2002; Ågren et al. 2013). In a survey conducted in 2001, the proportion of the short-scaped morph varied from zero to 100% (median 19 %, N = 69 populations; Ågren et al. 2013). The long-scaped morph is better pollinated, but also subject to more intense seed predation (Ehrlén et al. 2002; Ågren et al. 2006) and grazing damage compared to the short-scaped morph (Ågren et al. 2013). Seeds are dispersed from late June to late July, and may either germinate and give rise to seedlings the following year or enter the seed bank for one or more years (Toräng et al. 2010). Seedlings form a small rosette, and typically do not flower during their first year.

#### Study sites and data collection

This study used an experimental setup that was established in 2004 to assess the influence of ungulate grazers on selection regime and evolution of *Primula farinosa* populations (Ågren et al. 2013). In nine populations, two permanent plots were established and randomly assigned to either a control or a fenced exclosure (120 cm tall, 15 cm x 15 cm mesh size fence). Control and exclosure plots area varied between 70 m^2^ and 960 m^2^ depending on plant density and shape of the local population. During the first six years of the study, vegetation rapidly grew taller in exclosures, but from 2010 and onwards the difference in vegetation height between exclosures and control plots have stabilized (Thomann et al. 2018).

Using this experimental system, we conducted a demographic study from 2010 to 2016. Depending on population density, we established a minimum of 4 permanent quadrats (1m x 1m) in each control and exclosure plot. The number of plots was set to include ≥125 individuals in each population × treatment combination at the start of the experiment. We mapped all *Primula farinosa* individuals found within each quadrat, including every new individual appearing during the experiment. Each year from 2010 to 2016, we recorded survival, rosette diameter, and flowering status (vegetative or flower-producing) of each plant during flowering in May. At the time of fruit maturation in early July, we noted for each flower-producing plant whether the inflorescence had been removed by herbivores, the total number of flowers produced, the number of flowers that had initiated fruit development (a measure of pollination success; Ehrlén et al. 2002, Vanhoenacker et al. 2006), the number of initiated fruits consumed by the seed predator, the number of smut-infected flowers, and the number of intact fruits produced. To quantify plant size, rosette diameter was measured with a caliper to the nearest millimeter. The mean frequency of the short-scaped morph in the study quadrats in 2010-2015 was 0.46 (range 0.27-0.61, N = 9 populations) in the control and 0.38 (0.18-0.51) in the exclosures (Supplemental Table 1).

Grazed individuals were identified as those plants where the inflorescence was cut off but the base of the scape still visible, and for each plot grazing intensity was quantified as the proportion of flowering individuals grazed. The fences effectively keep ungulate grazers out, but do not prevent rodents or slugs from reaching plants inside the exclosures. Grazing damage and vegetation height did not show any temporal trend in control plots between 2005 and 2016 (Thomann et al. 2018). In contrast, damage from small herbivores in the exclosures was very low in the first years of the experiment (< 1% in 2005-2006), but increased over time as the vegetation grew taller in the exclosures (see Supplement of Thomann et al. 2018). This is consistent with the idea that increased vegetation cover and litter accumulation in exclosures increase the abundance of small herbivores. During the present study, grazing from small herbivores affected on average 4% of flowering plants in exclosures. In calculations below, we assume that this type of grazing does not affect surrounding areas to any appreciable extent and that the contribution of small mammals and slugs to damage in control plots is negligible.

### Statistical Analyses

#### Vital rates and intensities of biotic interactions

We used mixed-model ANOVA to examine the effects of treatment (exclosure vs. control; fixed factor), and year, site, and the treatment × site interaction (random factors) on recruitment rate (number of seedlings per seed produced), survival, rosette size, proportion of established plants flowering, proportion of flower-producing plants that were grazed, number of flowers produced by reproductive plants, proportion of flowers initiating fruit development, proportion of initiated fruits being consumed by the seed predator, and the number of mature intact fruits produced by flowering plants. The analyses were based on yearly treatment × site means.

#### Integral projection models (IPM)

To estimate population growth rates in control and exclosure plots in each population, we used integral projection models (IPMs) based on vital rate data collected between 2010 and 2016. IPMs allow modelling of population dynamics for species where vital rates such as survival and fecundity depend on a continuous predictor, such as individual size, but also allow explicit modelling of discrete life cycle stages such as seed banks (Ellner & Rees 2006; Merow et al. 2014). We constructed IPM functions as a combination of discrete stages, which describe population dynamics within a seed bank (Equation 1) and a seedling cohort (Equation 2), and a continuous size-structured stage representing the established plants (Equation 3). Equation (1) is the function for number of seeds in the seed bank (*n*_*SB*_), equation (2) is the function for number of seedlings (*n*_*SDL*_), and equation (3) is the state transition function for established plants (*n*_*E*_)

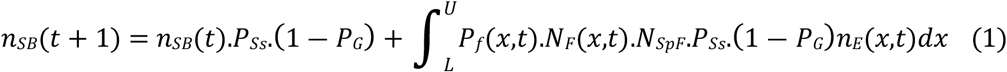

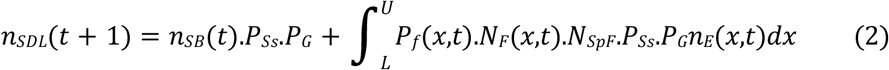

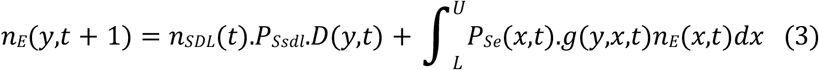

*L* and *U* represent the lower and upper limits of possible size (rosette diameter) in the size-structured elements of the IPM and were set as the minimum and maximum value observed for each site × treatment combination during the study, *x* represents size in year *t* and *y* represents size in year *t* + 1. Specific transitions are described as follows: *P*_*Ss*_ is the probability of a seed surviving, *P*_*G*_ is the probability of a surviving seed germinating, *P*_*Ssdl*_ is the probability of a seedling surviving, *P*_*f*_(*x,t*) is the probability of a established plant of size *x* to flower in year *t, N*_*F*_(*x,t*) is the expected number of intact fruits produced by a flowering plant of size *x* in year *t, N*_*SpF*_ is the number of seeds per fruit, *P*_*Se*_(*x,t*) is the probability of a established plant of size *x* to survive from year *t* to year *t* + 1, and *g*(*y,x,t*) is the probability of a size *x* established plant in year *t* to be of size *y* in year *t+1. D*(*y,t*) is the size distribution of newly established plants. We combined these three equations to create a transition kernel describing the contribution of vital rates to changes in population size and structure between years (following Rees et al. 2014). The dominant eigenvalue of the transition kernel then represents population growth rate (Ellner & Rees 2006; Rees et al. 2014).

In the case that the model assigned individuals sizes outside the upper and lower integration limits in the IPMs, we assigned them to a size corresponding to the nearest size limit (Merow et al. 2014). Moreover, as the relationship between size and fruit production was estimated using non-linear models, extrapolating outside the range of observed data under which the regression parameters were estimated can produce unrealistic predictions. In simulations, we therefore restricted the maximum predicted number of intact fruits produced to the observed maximum in the data from each site × treatment × year combination.

#### Vital rates for characterizing IPMs

To model the relationships between plant size and vital rates included in eqn. (3), vital rates were modeled as functions of rosette size separately for each site × treatment × year combination (cf. Merow et al. 2014). Vital rate functions were parameterized using generalized linear models, specifying logistic regression for survival between year *t* and year *t + 1*, and flowering status in year *t* (flowering vs. not flowering), least-squares regression for rosette size in year *t + 1*(growth), and Poisson regression for number of fruits produced by flowering plants in year *t*. In the IPMs, we retained all slope estimates irrespective of statistical significance, rather than assuming zero slopes for non-significant relationships.

Estimates of number of seeds per fruit, seed survival and probability of germination were obtained from previous demographic studies of *P. farinosa*. Number of seeds per intact mature fruit was set to 49.9, which was the mean obtained for 80 flowering plants in each of 16 populations on Öland sampled in two years (von Euler et al. 2014). Young *P. farinosa* seedlings are easy to miss at the census of the demographic plots in June, and recruits were therefore recorded only the year after germination. To estimate the size of the seed bank and the number of first-year seedlings, we followed the procedure outlined in Toräng et al. (2010). For each year, we estimated the number of first-year seedlings as the seed pool the previous year (number of seeds produced and number of seeds alive in the seed bank) multiplied by the estimate of germination probability (0.003883) obtained by Toräng et al. (2010). We then calculated seedling survival from the first to the second year by dividing the number of recruits observed the year after germination with the estimated number of first-year seedlings in the previous year.

#### Population growth rate (λ)

We calculated population growth (λ) estimates from IPMs for each annual transition, and then used this output to estimate stochastic growth rates (λs) for each site and treatment over the 2010-2016 period. Stochastic growth rates were calculated using the stoch.growth.rate function of the popbio package in R (Stubben et al. 2007) across the 6 yearly transitions, using default parameters. For Olstorp 5, we restricted the analyses to 5 transitions as the number of plants in the exclosure had become very small during the 2015/2016 transition.

#### Estimating direct and indirect effects of ungulate grazing on population dynamics

For each population, we quantified the direct, indirect, and net effects of ungulate grazers on population growth rate through three pairwise comparisons. We quantified the direct effect of ungulate grazers as the difference in stochastic lambda obtained by a model including all observations in the control treatment and a model that included only individuals in the control not damaged by grazers (Δλ_dir_ = λ_C_all_ – λ_C_undam_). The indirect effect of ungulate grazers on population growth rate was quantified as the difference between the stochastic lambda estimated for individuals in the control not damaged by grazers and that estimated for all individuals in the exclosure (Δλ_indir_ = λ_C_undam_ – λ_Excl_). Finally, the net effect of ungulate grazers was estimated as the difference between stochastic lambda calculated based on all plants in the control and that based on all plants in the exclosure (Δλ_net_ = λ_C_all_ – λ_Excl_ = Δλ_dir_ + Δλ_indir_).

#### Distinguishing indirect effects due to interactions with pollinators, seed predators, and small herbivores

We used simulations to estimate indirect of effects of grazers on population growth due to effects on interactions with pollinators, seed predators, and small herbivores. To examine the extent to which pollen availability contributed to differences in population growth rate between exclosures and control plots, we compared for each population and year the estimate of population growth rate in the exclosure based on observed vital rates only, with that obtained using models where the observed fruit initiation of each plant had been replaced by the average fruit initiation in the control plot. Similarly, to estimate the effects due to seed predation, we compared estimates of population growth rate based on observed vital rates only, with that obtained using models where the proportion of initiated fruits consumed by the seed predator had been replaced by the average proportion observed in the control plot. To determine the extent to which ungulate grazers affect *P. farinosa* population growth rate by reducing damage by small herbivores, we calculated for each exclosure the difference between the estimate of stochastic lambda obtained based on ungrazed plants only and that obtained based on all individuals. This difference provides an upper estimate of the indirect effects of ungulate grazers due to interactions with small herbivores since it assumes that no damage from small herbivores occurs in the control plots (see above).

#### Life Table Response Experiment

We used Life Table Response Experiment (LTRE) to examine how differences in establishment (the sum of the matrix elements representing the transition from seed to established plants), survival/growth (the sum of the matrix elements representing survival and growth of established plants), and reproduction (the sum of the matrix elements representing the number of seeds produced) contributed to differences in population growth rate (Caswell 1989, 2010).

For each treatment × population combination, a deterministic projection matrix was estimated as the mean matrix of annual projection matrices. We then calculated LTRE contributions as:

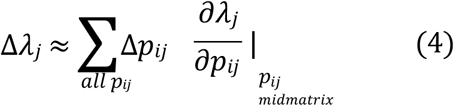

where Δ*λ*_*j*_ is the difference in deterministic growth rate between exclosure and control at site j, Δ*p*_*ij*_ is the difference in the value of the matrix element *i* between exclosure and control at site j, and 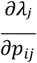 is the sensitivity of population growth rate to variation in matrix element i, evaluated with the midpoint matrix between exclosure and control of site *j*.

## RESULTS

### Differences in vital rates between treatments

Ungulate grazers were responsible for the great majority of grazing damage observed, and thereby directly and negatively affected plant fitness. The proportion of flowering plants whose inflorescence was grazed was on average six times higher in control plots compared to exclosures (least-square means, 24% vs. 4%; Table 1, Fig. 1E).

**Table 1.** Effect of treatment (exclosure vs. control) on nine fitness components: number of recruits per seed, survival, size (rosette diameter), proportion of plants flowering, proportion of flower-producing plants with inflorescence removed by grazers, number of flowers per flower-producing plant, proportion of flowers initiating fruit development, proportion of initiated fruits consumed by the seed predator, and number of intact fruits formed on flower-producing plants. The effect of treatment was tested with a mixed model, which also included year, population, and the population × treatment interaction as random factors. Denominator degrees of freedom (df) are given, and statistically significant effects are indicated in bold.

| Variable | df | F | P |
| --- | --- | --- | --- |
| Recruits per seed | 8 | 22.50 | **0.0015** |
| Survival | 8 | 0.084 | 0.7792 |
| Rosette diameter (mm) | 8 | 70.04 | **<0.0001** |
| Proportion flowering | 8 | 0.060 | 0.8126 |
| Proportion grazed | 7.75 | 23.06 | **0.0015** |
| Number of flowers | 8.14 | 45.51 | **0.0001** |
| Fruit initiation | 8.02 | 2.80 | 0.1328 |
| Fruit predation | 7.97 | 1.58 | 0.2443 |
| Number of intact fruits | 7.70 | 2.32 | 0.1673 |

**Figure 1.**
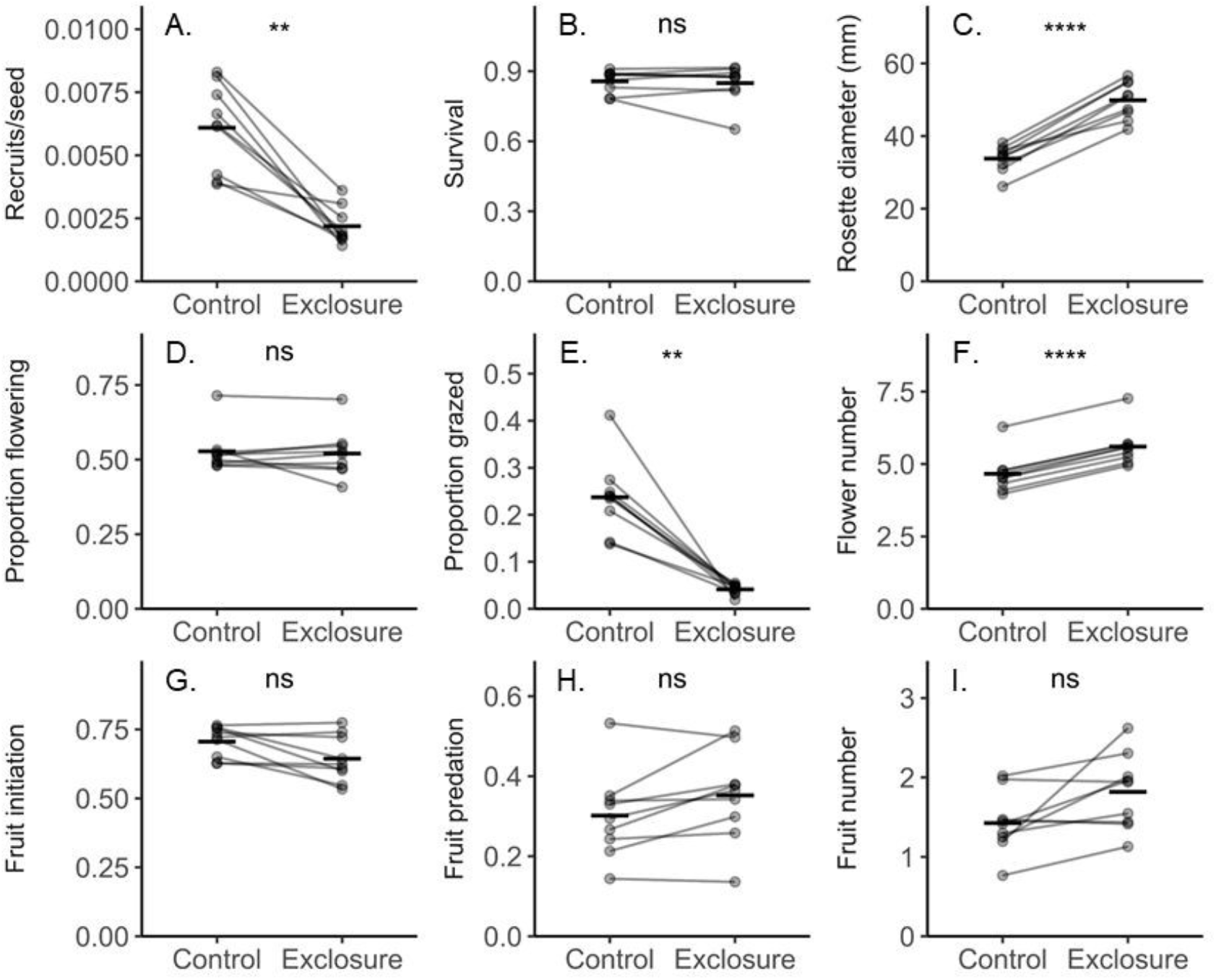
Least-square means of vital rates (number of recruits per seed, survival, rosette diameter, proportion of plants flowering, and number of flowers and number of intact fruits produced by flowering plants) and estimates of interactions with grazers, pollinators (quantified as fruit initiation), and seed predators (quantified as proportion of initiated fruits predated) in the control and in exclosures. Least-square means for each site × treatment combination (circles) and the grand treatment mean (horizontal bar) are indicated. Statistically significant effects of treatment were found for recruitment rate, rosette size, proportion of plants grazed, and number of flowers produced by flower-producing plants (Table 1).

Ungulate grazers also indirectly affected plant fitness. Exclusion of ungulate grazers reduced the recruitment rate of *P. farinosa* by on average 64% (least-square means, 0.00219 recruits per seed in exclosures vs. 0.00609 recruits per seed in the control; Fig. 1A), but was associated with a 48% increase in *P. farinosa* rosette diameter (49.9 vs. 33.8 mm; Fig. 1C), and a 20% increase in the average number of flowers (4.66 vs. 5.60; Fig. 1F; Table 1). Contrary to prediction, fruit initiation and fruit predation were not significantly higher in controls than in exclosures (Fig. 1G, H; Table 1). Differences in mean fruit initiation and fruit predation between control and exlosure were large in some populations, but the direction of effect varied. The difference in mean fruit initiation varied from 2% lower to 18% higher in the control compared to the exclosure (mean 6% higher, N = 9 populations) and estimates of fruit predation from 16% lower to 4% higher (mean 5% lower). Also survival of established plants, the proportion of established plants that flowered, and number of intact mature fruits produced by flowering plants were not significantly affected by ungulate grazers (Table 1; Fig. 1B, D, I).

### Effect of ungulate grazers on P. farinosa *population growth rate*

Direct effects, in terms of damage to inflorescences by ungulate grazers, reduced stochastic population growth rate, but this was more than balanced by positive indirect effects at 7 of the 9 sites (Fig. 2). Estimates of population growth rate of control plants with and without damaged plants included, indicated that the damage inflicted by ungulates to *P. farinosa* inflorescences reduced λs by on average 0.044 (paired t-test, P = 0.005). Comparison of population growth rates in exclosures and of undamaged plants in controls suggested that the indirect effect of ungulates was positive at all sites but two, and on average more than twice as large as the negative direct effects (Δλ_indirect_ = 0.101; paired t-test, P = 0.04), resulting in a positive net effect of ungulate grazing (Δλ_net_ = 0.058; paired t-test, P = 0.109; Fig. 2).

**Figure 2.**
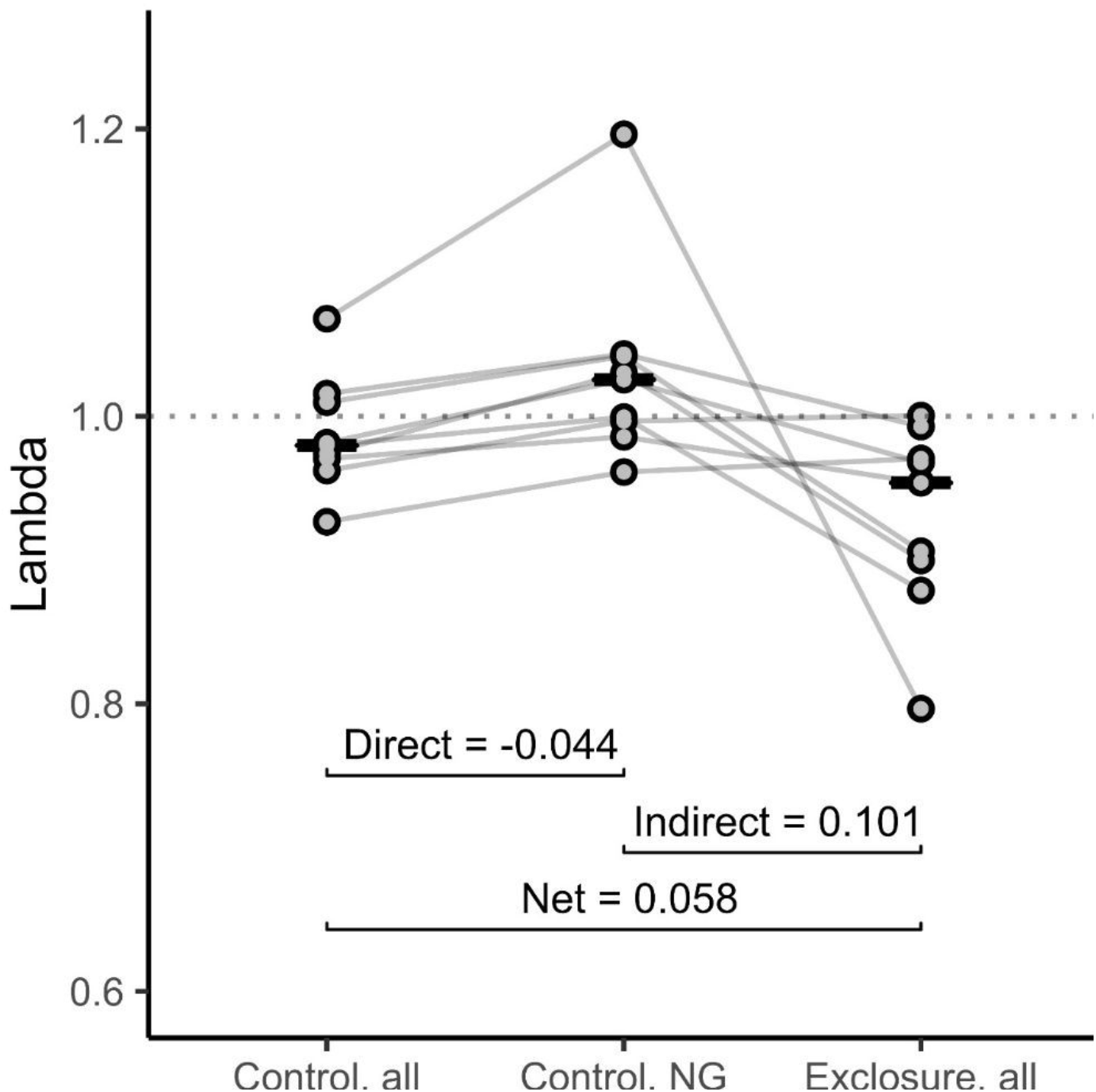
Stochastic lambda estimated for all plants in the control treatment (Control, all; λ_C_all_), for plants in the control escaping damage from grazers (Control, NG; λ_C_undam_), and for plants in exclosures (Exclosure, all; λ_Excl_). Lines connect symbols representing estimates for each of the 9 study sites, and horizontal bars represent mean values in each treatment. The direct effect of ungulate grazers was quantified as Δλ_dir_ = λ_C_all_ – λ_C_undam_, the indirect effect as Δλ_indir_ = λ_C_undam_ – λ_Excl_, and the net effect as Δλ_net_ = λ_C_all_ – λ_Excl_. Effect sizes shown are means of effects across sites. Statistical significance of effects based on paired t-tests: direct effect, P = 0.005, indirect effect, P = 0.04, net effect, P = 0.109.

### Causes of indirect effects

Simulations of the effects of differences in pollination intensity, seed predation and damage from small herbivores suggested that effects of ungulate grazing via these interactions were small (Fig. 3). At all sites, damage by small herbivores slightly reduced estimates of population growth rate in the exclosures, suggesting a weak positive indirect effect of ungulate grazers via this interaction (mean effect size = 0.01; Fig. 3).

**Figure 3.**
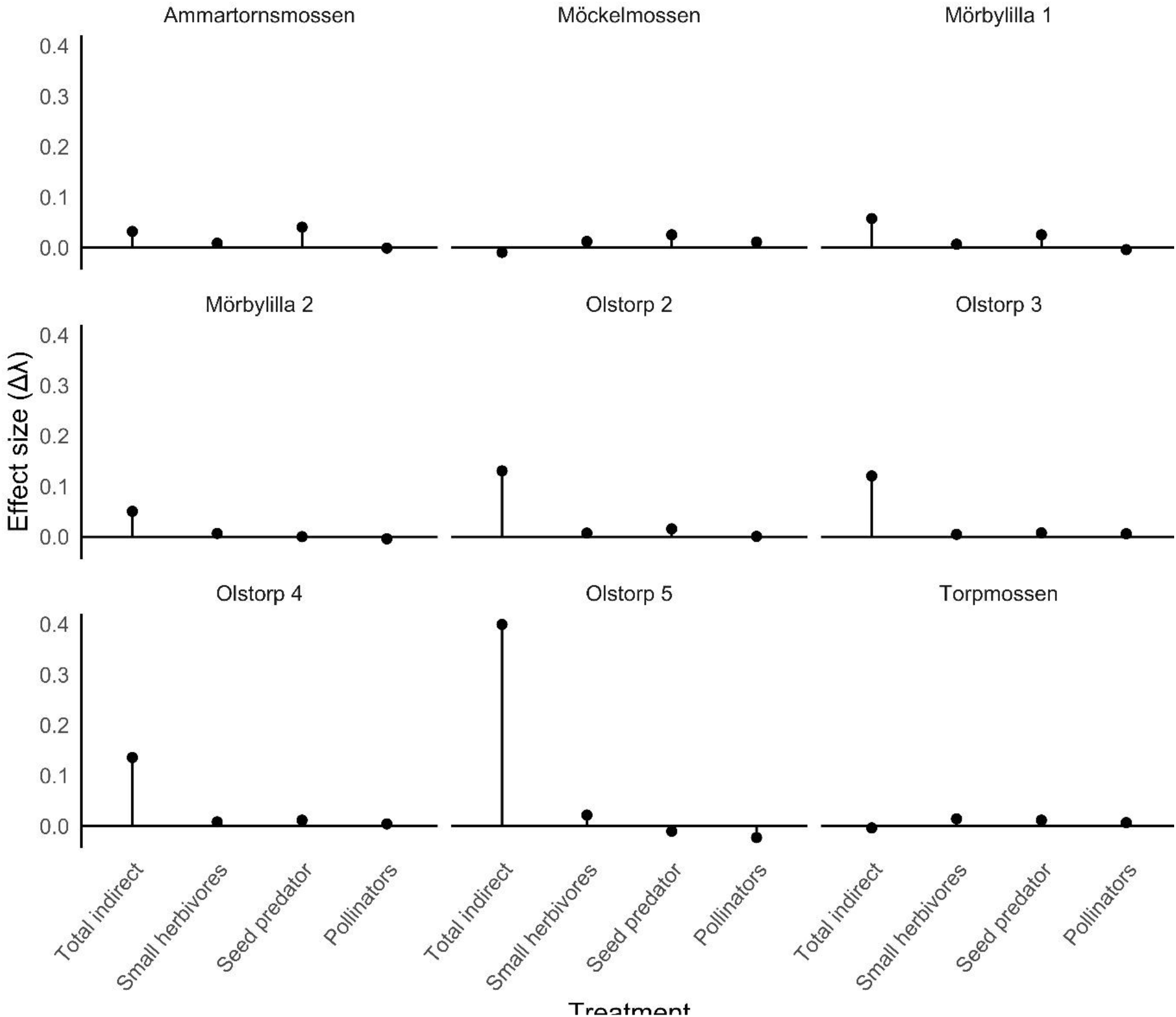
Indirect effect of ungulate grazers on *P. farinosa* population growth rate at nine sites on Öland, S. Sweden. The total indirect effect is indicated as well as contributions from effects via interactions with small herbivores, seed predators, and pollinators estimated with simulations (see Methods).

### Effects of exclosure on different life history stages (LTRE)

LTRE analyses indicated that lower establishment could explain much of the reduced population growth rate in exclosures compared to controls, and that lower establishment was in contrast to higher survival/growth in exclosures than in controls at several sites (Figure 4).

**Figure 4.**
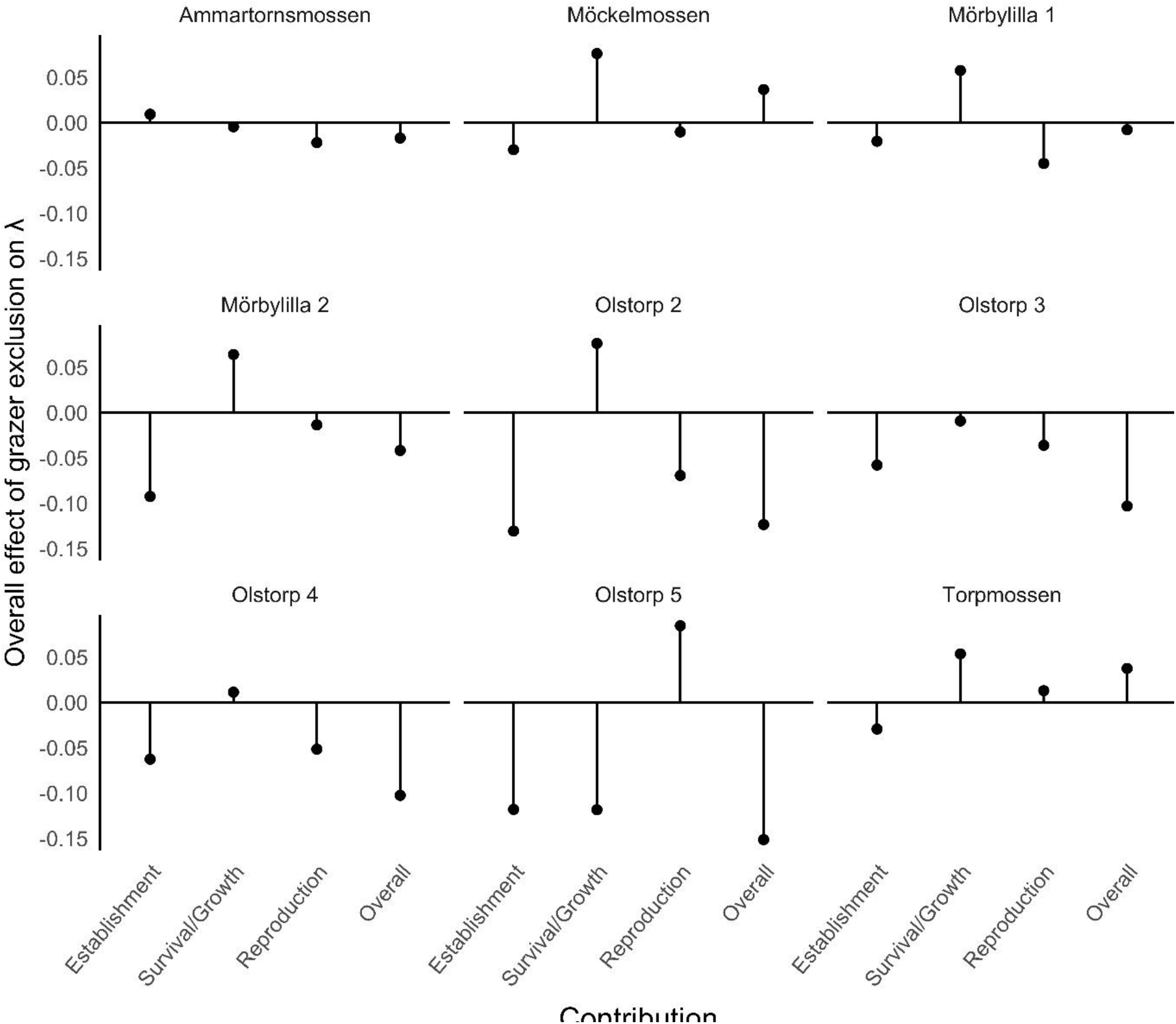
Life table response experiment (LTRE) by site. The individual and summed contributions of three main life history stages (establishment, survival/growth, and reproduction) to the difference in lambda between exclosure and control are indicated. Negative values indicate that exclusion of ungulate grazers reduces the performance of *P. farinosa*.

## DISCUSSION

This study shows that both direct and indirect effects of ungulate grazers influence population dynamics of the rosette-forming, grassland herb *P. farinosa*. We identified a negative direct effect of grazing damage on population growth rate, but this was more than balanced by a positive indirect effect, which was linked to increased recruitment in grazed areas. Simulations indicated that indirect effects via interactions with pollinators, seed predators, and small herbivores played a limited role for the positive indirect effects observed. Instead, we suggest that creation of gaps in the vegetation through trampling and reduced competition for light and nutrients cause most of the positive indirect effect.

Grazing of reproductive parts may reduce seed production in many systems (e.g., Root 1996; Gómez 2005), but the effects on population growth has seldom been assessed. Ungulate grazers do not selectively feed on *P. farinosa*. Instead, while feeding on surrounding vegetation they also consume inflorescences of *P. farinosa*, which thwarts seed production by the damaged individual and negatively affects population growth. In the present study, 24% of the flowering *P. farinosa* had their inflorescence removed by grazers in the control plots. This damage was estimated to reduce population growth rate by 0.044 on average across the nine populations. Typically, only the inflorescence was damaged, which may explain why the direct negative effect on the population growth rate of *P. farinosa* was not even stronger than that observed. Like other rosette-forming and prostrate herbs, *P. farinosa* avoids damage by ungulate grazers to vegetative parts better than do herbs with an upright growth form (Díaz et al. 2001; Fujita & Koda 2015).

Ungulate grazers positively affected the recruitment rate, and the LTRE analysis indicated that this could explain much of the overall positive indirect effect of grazers on *P. farinosa* population growth rate. This positive effect is likely to be related to the physical disturbance of the vegetation, and reduced resource competition with other plant species.

Through trampling the ungulate grazers create gaps in the ground vegetation, which should have contributed to the higher seedling establishment observed in control plots than in exclosures in all but one population (Figs. 1A, 4). Moreover, by reducing vegetation height and the size of co-occurring plant species, the grazers reduce litter accumulation as well as competition for light and other resources. This should facilitate *P. farinosa* establishment, growth, and flowering: Although established plants on average were larger in exclosures, there was no significant difference between treatments in survival, probability of flowering, or in the number of intact fruits produced by flowering plants (Fig. 1 B, C, D, I). Increased seedling establishment after litter removal and disturbance of the vegetation has been recorded in several grassland species (e.g., Bullock et al. 1994; Jakobsson & Eriksson 2000; Wilsey & Polley 2003), and in some species, including the congener *P. veris*, this has been shown to strongly influence estimates of population growth rate (Ehrlén et al. 2005; Lemmer et al. 2021). This highlights the potentially strong indirect effects of grazing on population growth, and is consistent with the observation that intermediate levels of grazing or other disturbance is favourable for the persistence of species that depend on recruitment from seed.

Contrary to prediction, pollination success, quantified as fruit initiation, and the intensity of seed predation were not significantly lower in the exclosures compared to controls. Instead, the direction of the differences between exclosure and control varied among sites (Fig. 1G, H). Although differences were large at some sites (fruit initiation differed by up to 18% and fruit predation by up to 16%), simulations indicated that these differences had only small effects on stochastic population growth rate in all populations (Fig. 3). The results indicate that sensitivity of population growth rate to changes in seed production per flowering plant escaping grazing damage was low. At least two factors may have contributed to the absence of a significant overall effect of grazer exclusion on intensities of pollination and seed predation. First, following the establishment of the experiment, the frequency of the short morph has decreased in exclosures (Ågren et al. 2013). During the present study, the proportion of the short-scaped morph was lower in the exclosure compared to the control in all but two populations (Olstorp 4 and Torpmossen). This should have contributed to the relatively weak effects observed since both pollination intensity and seed predation are less affected by vegetation height in the long-scaped compared to the short-scaped morph (Thomann et al. 2018). Second, in all populations the mean number of flowers per inflorescence was greater in the exclosure than in the control (Fig. 1F). No association between number of flowers and fruit initiation was detected in an earlier study (Vanhoenacker et al. 2006). However, seed predation has been found to increase with number of flowers in *P. farinosa* (Vanhoenacker et al. 2009) suggesting that the greater number of flowers per flowering plant in exclosures may have contributed to that we did not find lower intensities of seed predation when grazers were excluded.

By reducing vegetation height and litter accumulation, ungulate grazers may reduce the impact of small herbivores such as rodents and slugs finding shelter from predators in tall vegetation (e.g., Goheen et al. 2010). In the first years following the establishment of the experiment (i.e., in the years preceding the present study), vegetation grew successively taller in the exclosures, and so did damage to *P. farinosa* inflorescences from small herbivores (Thomann et al. 2018). This suggests that ungulate grazers positively affect the fitness of *P. farinosa* by reducing damage from small herbivores. However, damage from small herbivores affected on average only 4% of all inflorescences in exclosures during the present study, and simulations indicated that the effects on stochastic population growth rate were weak (Fig. 3).

In summary, grazing may affect plant fitness and population dynamics through a multitude of mechanisms. Here we have shown that the negative direct effect of damage from ungulate grazers is more than balanced by positive indirect effects on population growth rate of the grassland herb *P. farinosa*. Plant species are often categorized as those favoured by grazing and those disfavoured by grazing, essentially corresponding to species that are more abundant in grazed areas than in non-grazed areas, and species for which the opposite is true. We suggest that the quantification of direct and indirect effects of grazers on plant population growth rate provides insight into the causes of this variation, and a better understanding of the effects of herbivores on plant fitness, abundance and distribution.

## ACKNOWLEDGEMENTS

We thank a large number of dedicated field assistants and students for help with data collection. This research was financially supported by grants from Formas and the Swedish Research Council to J. Ågren and J. Ehrlén.

## AUTHOR CONTRIBUTIONS

JÅ and JE designed the study; JÅ, JE, and MT collected data; TR and MT analysed data; TR wrote the first version of the manuscript; all authors contributed substantially to revisions.

## DATA AVAILABILITY STATEMENT

The data supporting the results will be deposited in the Dryad digital repository after acceptance of the paper.

---

**Supplemental Table 1.**
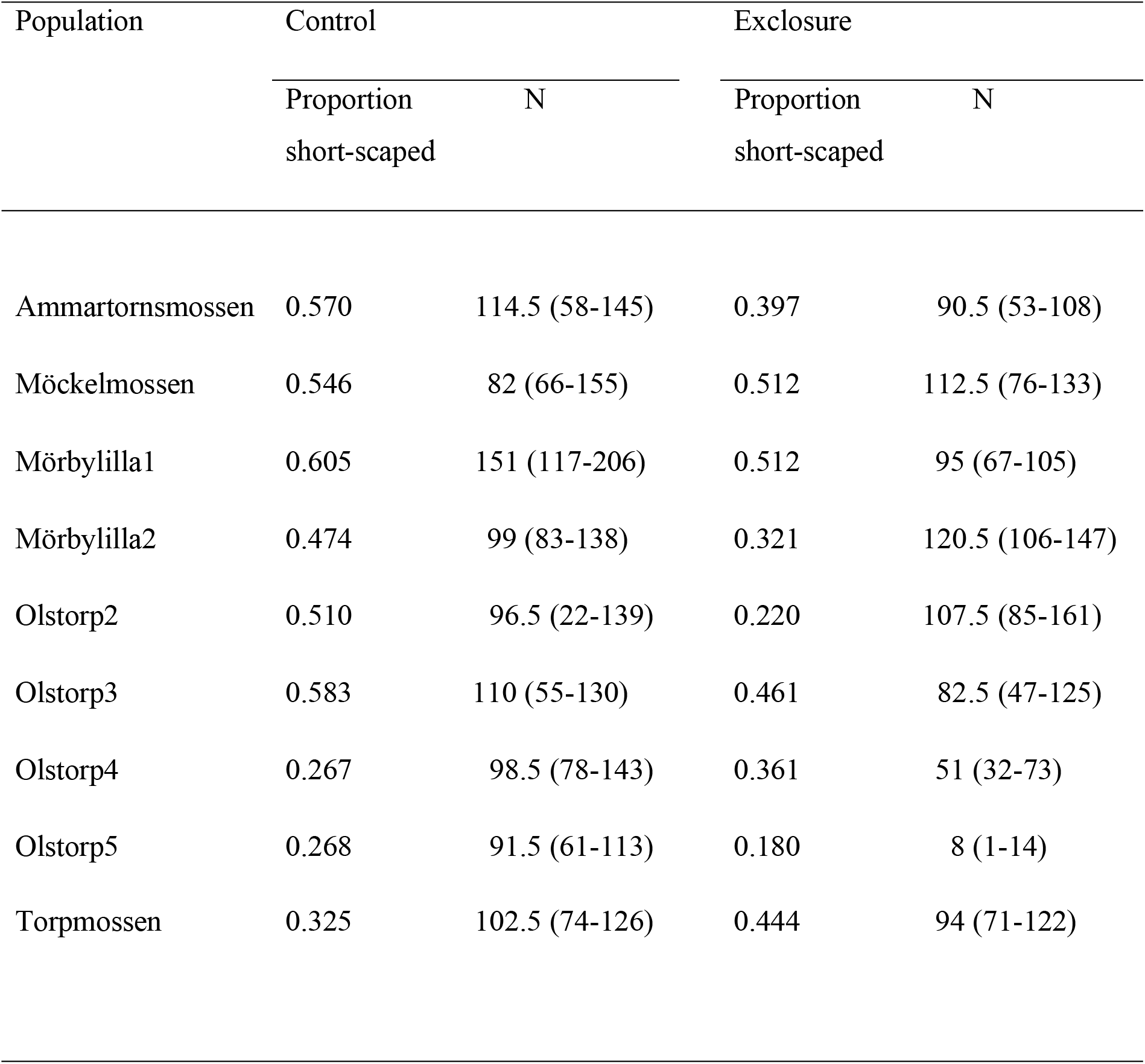
Average frequency of the short-scaped morph of *Primula farinosa* among flowering plants in the demographic quadrats in the control and in exclosures during the period 2010-2015 (N = 6 years, except Olstorp 5 Exclosure N =5). The median and range in sample size in individual years are given.

